# Task Activation Results in Regional ^13^C-Lactate Signal Increase in the Human Brain

**DOI:** 10.1101/2024.02.01.577808

**Authors:** Biranavan Uthayakumar, Nicole I.C. Cappelletto, Nadia D. Bragagnolo, Albert P. Chen, Nathan Ma, William J. Perks, Ruby Endre, Fred Tam, Simon J. Graham, Chris Heyn, Kayvan R. Keshari, Hany Soliman, Charles H. Cunningham

**Affiliations:** Department of Medical Biophysics, University of Toronto, Toronto, Ontario, Canada; Physical Sciences, Sunnybrook Research Institute, Toronto, Ontario, Canada; GE Healthcare, Toronto, Ontario, Canada; Pharmacy, Sunnybrook Health Sciences Centre, Toronto, ON, Canada; Radiology, Sunnybrook Health Sciences Centre, Toronto, ON Canada; Department of Radiology, Memorial Sloan Kettering Cancer Center, New York, NY, USA; Molecular Pharmacology Program, Memorial Sloan Kettering Cancer Center, New York, NY, USA; Radiation Oncology, Sunnybrook Health Sciences Centre, Toronto, ON, Canada

## Abstract

Hyperpolarized-^13^C magnetic resonance imaging (HP-^13^C MRI) was used to image changes in ^13^C-lactate signal during a visual stimulus condition in comparison to an eyes-closed control condition. Whole-brain ^13^C-pyruvate, ^13^C-lactate and ^13^C-bicarbonate production was imaged in healthy volunteers (N=6, ages 24-33) for the two conditions using two separate hyperpolarized ^13^C-pyruvate injections. BOLD-fMRI scans were used to delineate regions of functional activation. ^13^C-metabolite signal was normalized by ^13^C-metabolite signal from the brainstem and the percentage change in ^13^C-metabolite signal conditions was calculated. A one-way Wilcoxon signed-rank test showed a significant increase in ^13^C-lactate in regions of activation when compared to the remainder of the brain (*p* = 0.02, V = 21). No significant increase was observed in ^13^C-pyruvate (*p* = 0.11, V = 17) or ^13^C-bicarbonate (*p* = 0.95, V = 3) signal. The results show an increase in ^13^C-lactate production in the activated region that is measurable with HP-^13^C MRI.

## Introduction

Hyperpolarized-^13^C magnetic resonance imaging (HP-^13^C MRI) is a minimally invasive technique that enables imaging of injected ^13^C-labelled metabolites and their downstream conversion products *in vivo*. The most commonly imaged metabolites are ^13^C-pyruvate and its downstream products ^13^C-lactate and ^13^C-bicarbonate. The production of ^13^C-lactate signal indicates the reduction of ^13^C-pyruvate to ^13^C-lactate in the cytosol, while ^13^C-bicarbonate production indicates the decarboxylation of ^13^C-pyruvate into acetyl-CoA on the mitochondrial membrane. HP-^13^C MRI has previously been used to image ^13^C-metabolite production in the human brain in healthy and disease states (2, 3, 4, 5, 6, 7).

Previous results showed concordant regional patterns of ^13^C-pyruvate, ^13^C-lactate and ^13^C-bicarbonate across individuals suggesting a highly regulated process leading to the pattern (5). At the same time, increased lactate concentration in the brain has been measured using proton spectroscopy and a visual stimulus (9, 10, 11). However, the effect of a visual stimulus, and the concomitant increase in lactate pool size, on the ^13^C-metabolite pattern in the brain has not been documented previously. We hypothesized that the ^13^C-metabolite distribution detected with HP-^13^C MRI would be modulated in response to task activation. In this study whole-brain HP-^13^C MRI was used to assess regional changes in ^13^C-lactate signal during a visual task stimulus. Blood-oxygen level dependent fMRI (BOLD-fMRI) images were acquired to delineate regions of BOLD activation in each participant. We hypothesized that the BOLD-fMRI defined regions of activation would have higher ^13^C- lactate signal under stimulus conditions relative to control conditions.

## Materials and Methods

Written informed consent was obtained from six study participants (3 male, 3 female) and procedures were performed under a protocol approved by the Research Ethics Board of Sunnybrook Health Sciences Centre and by Health Canada under a Clinical Trial Application in compliance with the ICH-GCP and Declaration of Helsinki. Participants ranging in age from 24 to 33 (mean age of 26) were screened for cognitive impairment using the Montreal Cognitive Assessment (12).

Several hours prior to scanning, two doses of ^13^C-pyruvic acid were prepared using a 1.47 g sample of [1-^13^C]pyruvic acid (Sigma Aldrich, St. Louis, MO) for each dose and polarized in a SPINLab polarizer system (GE Healthcare, Waukesha, WI) for a minimum of two hours. Immediately prior to scanning, a 20 gauge intravenous catheter was placed into the forearm of each participant before positioning them in the scanner bore.

Images were acquired using a GE MR750 3.0T MRI scanner (GE Healthcare, Waukesha, WI) using a ^13^C birdcage head coil built in-house for HP^13^C imaging. Volumetric images of ^13^C-lactate, ^13^C-bicarbonate, and ^13^C-pyruvate were acquired using a 3D echo-planar sequence (13) over a 60 s acquisition. Each metabolite had 24 spectrally-selective excitation pulses applied for the 24 slice direction phase encodes. The flip angle used for ^13^C-lactate and ^13^C-bicarbonate was 20^*°*^, while 2^*°*^ was used for^13^C-pyruvate, resulting in effective (net) flip angles of 80^*°*^ and 11^*°*^, respectively, for the acquisition of each volume. This was repeated every 5 s, giving a temporal resolution of 5 s with 15 mm isotropic spatial resolution covering the entire brain.

Each participant underwent two HP^13^C scans, consisting of a task (^13^C-task) and control (^13^C-control) scan. The visual task consisted of an 8 Hz flashing checkerboard stimulus initiated 5 s prior to the ^13^C-task scan. For the ^13^C-control scan, the participant was asked to close their eyes (and a blank screen with a plus sign was projected onto the screen). To control for any effect of scan ordering, three of the participants had the control scan first, whereas the other three had the control scan second. All stimuli and visual prompts were delivered via a projector and projector screen setup. The projector was placed inside the MRI console room facing into the MRI room, where a projector screen was setup. Prior to imaging, the projector was lined up with the projector screen and we verified with the participant in the scanner that they were able to view a test image. There was a half-hour wait period between ^13^C-task and ^13^C-control scans.

During the half-hour wait period, the ^13^C head coil was interchanged with an 8-channel ^1^H head coil (Invivo Inc, Pewaukee, WI) to perform an anatomical T1-weighted and BOLD fMRI scans. Anatomical T1-weighted images were acquired with an axial RF-spoiled gradient echo sequence (FOV 25.6×25.6 cm^2^, 1 mm isotropic resolution, repetition time (TR) = 7.6 ms, echo time (TE) = 2.9 ms, flip angle 11^*o*^). BOLD fMRI data were acquired using a single-shot EPI acquisition with 3.5 mm isotropic resolution, TR = 2000 ms, TE = 29 ms, 40^*o*^ flip angle, 64 by 64 matrix size and 38 3.5 mm slices with 0 mm gap. The task block duration was the same as the HP^13^C scan duration (one minute) and the total fMRI scan duration was 5 minutes, including 3 task blocks interleaved with 2 control blocks. The timing of the scan protocol is shown in Fig. 1.

**Figure 1.**
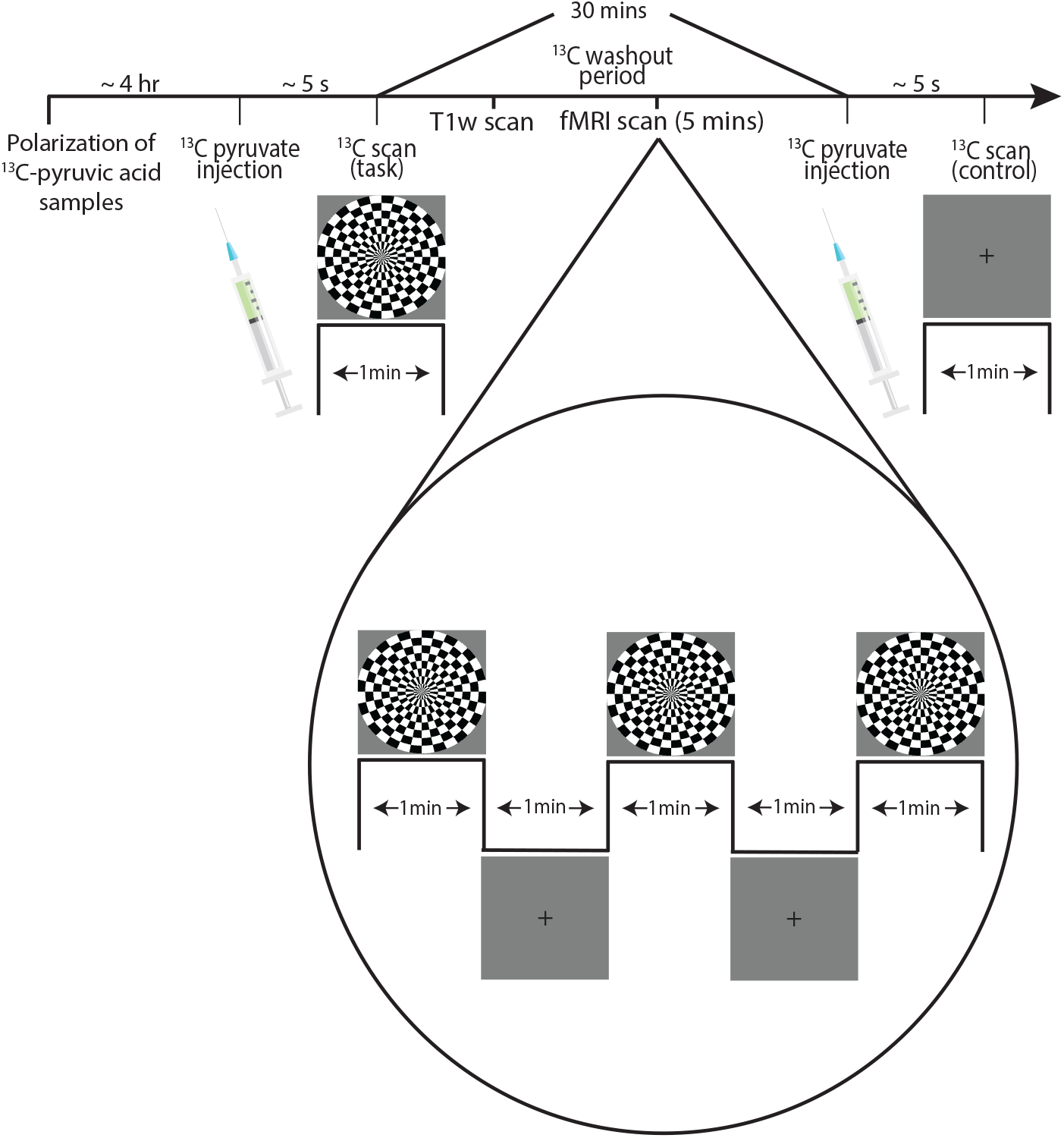
Image acquisition workflow for the visual stimulus scans. Two sets of ^13^C-images were acquired with a thirty minute wait period between acquisitions to allow for washout of residual ^13^C-metabolites. During this wait period, anatomical images and BOLD-fMRI images were acquired.

The ^13^C-metabolite images were summed across time to calculate the total ^13^C-metabolite signal during the 1 minute acquisition window. The ^13^C-lactate images were affine registered to the anatomical T1-weighted images using FSL (14), and the same transformation was applied to the ^13^C-pyruvate and ^13^C-bicarbonate images.

The T1-weighted anatomical images were parcellated into 132 brain regions according to the BrainCOLOR labelling protocol (15) using an automated workflow (16). These parcellation maps were then used to compute regional ^13^C-pyruvate, ^13^C-lactate, and ^13^C-bicarbonate signals for each participant using the mri segstats function (17). Regions with volumes below the voxel volume of 3.4 mL were excluded, resulting in 90 parcellated regions for each subject.

BOLD-fMRI pre- and post-processing was done using the Analysis of Functional Images (AFNI) toolbox (19, 20), and included removal of the first two TRs of each acquisition, registration across time to mitigate motion effects, and blurring with a Gaussian function (4mm FWHM). Voxelwise linear regression coefficients were fit using the stimulus block design as an independent variable, input as a repeating boxcar function with box length one minute. T-statistics were calculated for each positive regression coefficient using an alpha of 0.001. Cluster correction was used to control for multiple comparisons, with voxels in face-to-face contact comprising each cluster (21).

The BOLD-fMRI t-statistic map was used to define an an activation volume for each subject, with the remainder of the brain (including the white matter and ventricles) defined as the non-activation volume.

Each regional ^13^C-metabolite signal, as well as the signal from the activation volume and non-activation volume, were normalized by the ^13^C-metabolite signal from the brainstem region from the particular participant (referred to as ’brainstem normalization’ below). This was done to normalize for any global signal scaling between scans, such as those caused by differing ^13^C-pyruvate polarization levels, while maintaining the inter-regional pattern.

The percentage change in ^13^C-metabolite signal between task and control conditions was calculated as follows:

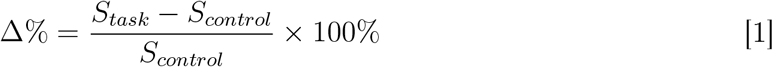

where *S*_*task*_ is the brainstem-normalized ^13^C-metabolite signal from a particular region/subject in the task condition and *S*_*control*_ is the signal from the same metabolite/region/subject in the control condition (also brainstem normalized). A one-way Wilcoxon signed-rank test was used to test for a difference in the Δ% between the activation and non-activation volumes.

The thresholded t-statistic maps from the BOLD fMRI scans were also used to designate parcellated brain regions as activated regions or non-activated regsions. The remaining parcellated brain regions were categorized as ’non-activation regions’.

## Results

Example t-statistic maps registered to the MNI152 atlas (22)) are shown in Fig. 2. Example ^13^C-task and control scans are shown in Fig.3. BOLD-fMRI activation was consistently achieved in the visual cortex over the course of the 5 minute block design in all six participants. This confirmed that the stimulus block length was sufficient for inducing activation during the HP^13^C scans within the one minute acquisition time. The Δ% for the activation and non-activation volumes is plotted in Fig.4.

**Figure 2.**
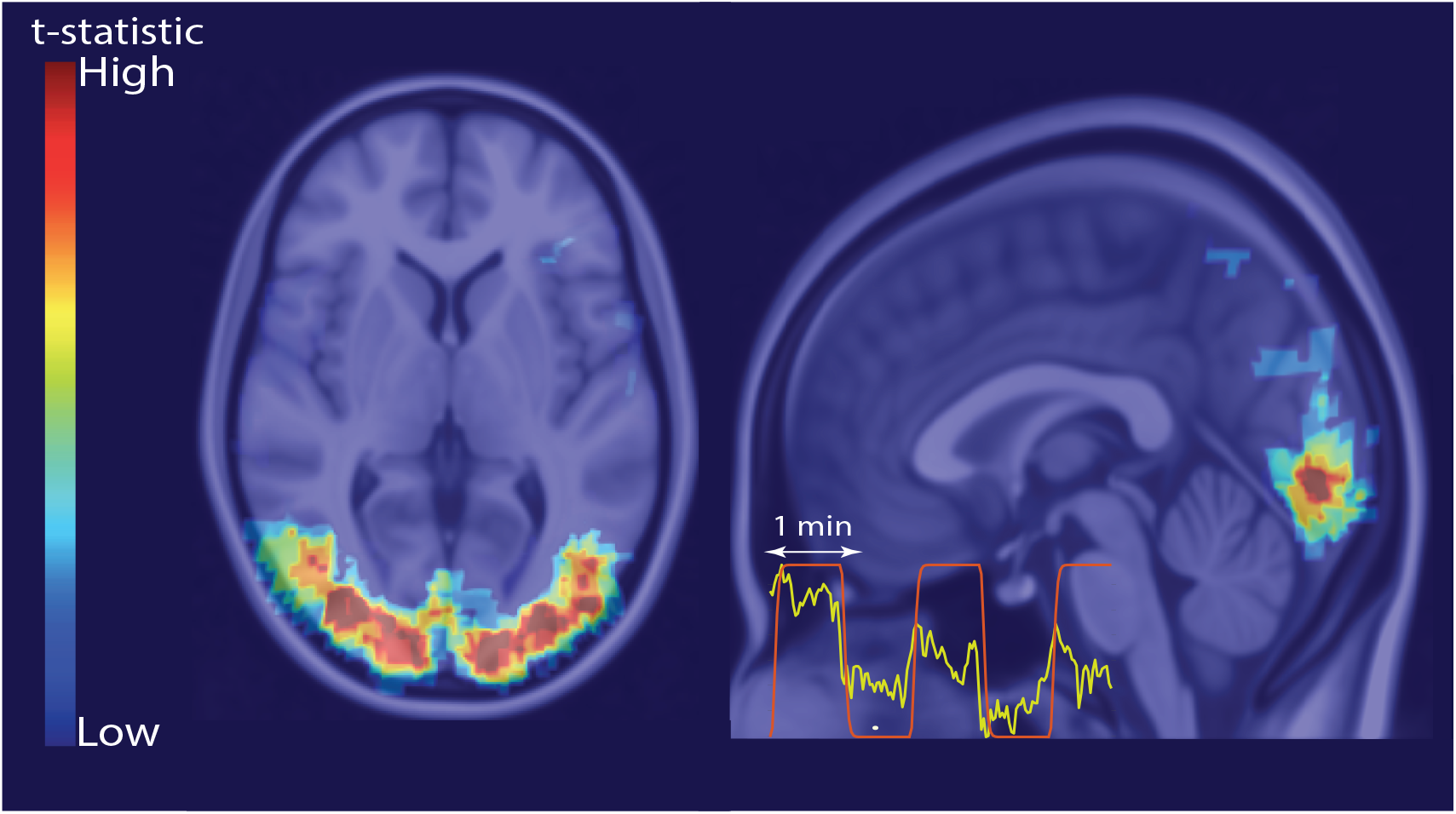
Example BOLD fMRI t-statistic map for the visual stimulus registered to the MNI152 atlas. The inset is an example time series curve from a voxel with significant activation (yellow) and the stimulus regressor used in the general linear model (red).

**Figure 3.**
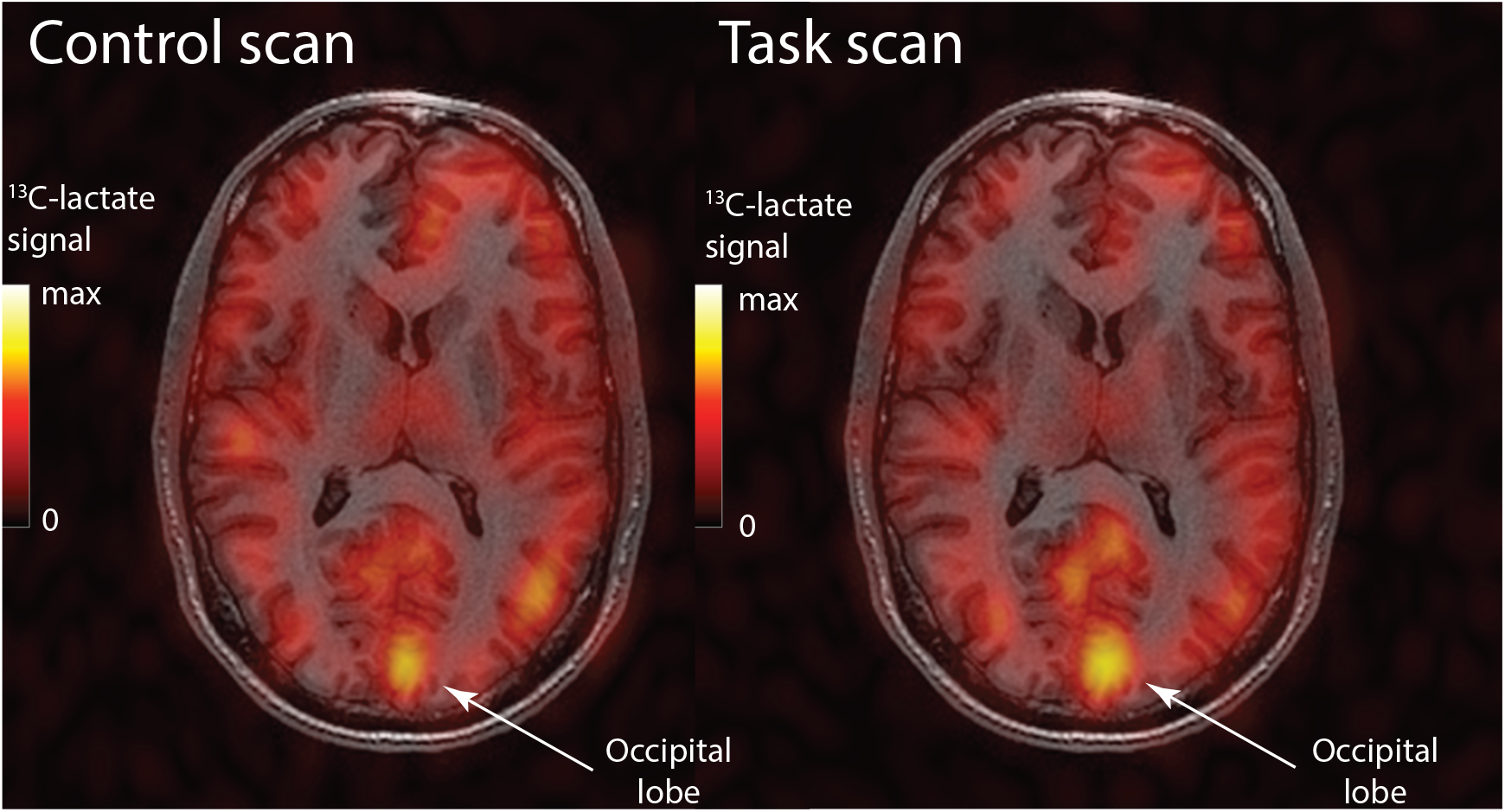
Example ^13^C-lactate images overlaid on the anatomical image for both the control and task conditions. Increases in ^13^C-lactate signal can be observed in the parts of the occipital lobe pointed out with the arrows. The same color bar scale was used for both images.

The one-way Wilcoxon signed-rank test showed a significant difference in Δ% for ^13^C- lactate signal between the activation and non-activation non-activation volumes (*p* = 0.02, V = 21), indicating a task-induced increase in ^13^C-lactate production in the activation volume relative to the rest of the brain. The Wilcoxon signed-rank tests for Δ% ^13^C-bicarbonate and Δ% ^13^C-pyruvate showed no significant difference between the activation and non-activation volumes (*p* = 0.95, V = 3 and *p* = 0.11, V = 17 respectively).

The parcellated regions withing the activation volume for all six individuals were: the bilateral calcarine cortex, cuneus, inferior occipital gyrus, lingual gyrus, middle occipital gyrus, occipital pole, occipital fusiform gyrus, and superior occipital gyrus. These regions are highlighted in Fig.5, with the Δ% ^13^C-lactate signal visibly higher in these regions in comparison to the non-activation volume regions.

## Discussion

Increased ^13^C-lactate signal in visual cortex regions was observed following a visual stimulus, consistent with prior studies using proton spectroscopy (23, 10, 24). These results demonstrate that task activation causes an increase in regional ^13^C-lactate signal measured with HP-^13^C MRI. Elevated regional neuronal energy needs are met through the hemodynamic response. However, the observed regional increases in ^13^C-lactate are unlikely to be due to a hemodynamic response-driven increase in vascular ^13^C-lactate, since the brain ^13^C-lactate signal has been shown to be largely parenchymal (25). Roughly 25% of astrocytic glutamate is released as lactate through the TCA cycle (26). Glutamate, the most abundant excitatory neurotransmitter in the brain, has consistently been observed to increase under stimulus conditions when measured through fMRS (27). Glutamate is cleared from the intracellular space by astrocytes through a process that is bioenergetically supported by the non-oxidative metabolism of pyruvate into lactate (28). However, lactate can also be used to meet transient energy needs in neurons (8), and neuronal stimulus directly induces neuronal lactate production (29). Taking these all into account, the working hypothesis is that the task-induced increase in ^13^C-lactate is due to a combination of astrocytic glutamate clearance rapidly meeting the energy demand of neuronal ring.

No significant differences between task and control conditions were detected for the ^13^C- pyruvate signal, although an increasing trend can be observed in Fig. 4. Increased cerebral blood volume due to functional activation would be expected to increase regional ^13^C- pyruvate signal, especially as ^13^C-pyruvate signal is largely in the vascular compartment (25). However, cerebral blood volume increases only slightly during functional activation (30), so it is possible that the effect size was too small to discern a significant change in ^13^C-pyruvate signal with the small sample size.

**Figure 4.**
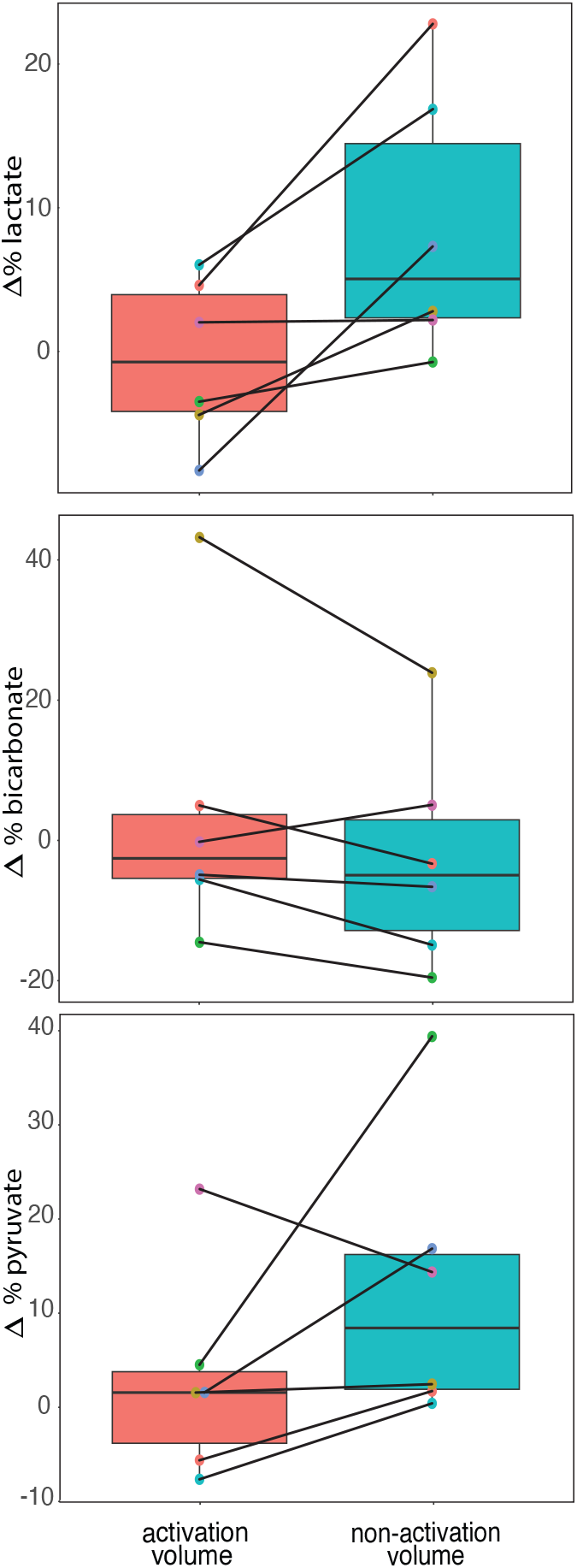
The percentage change between task and control ^13^C-scans for the non-activated volume (left) and activated volume (right). The lines connect the data points from the same individual, the box centre line indicates the mean and whiskers at the 25th and 75th percentile.

**Figure 5.**
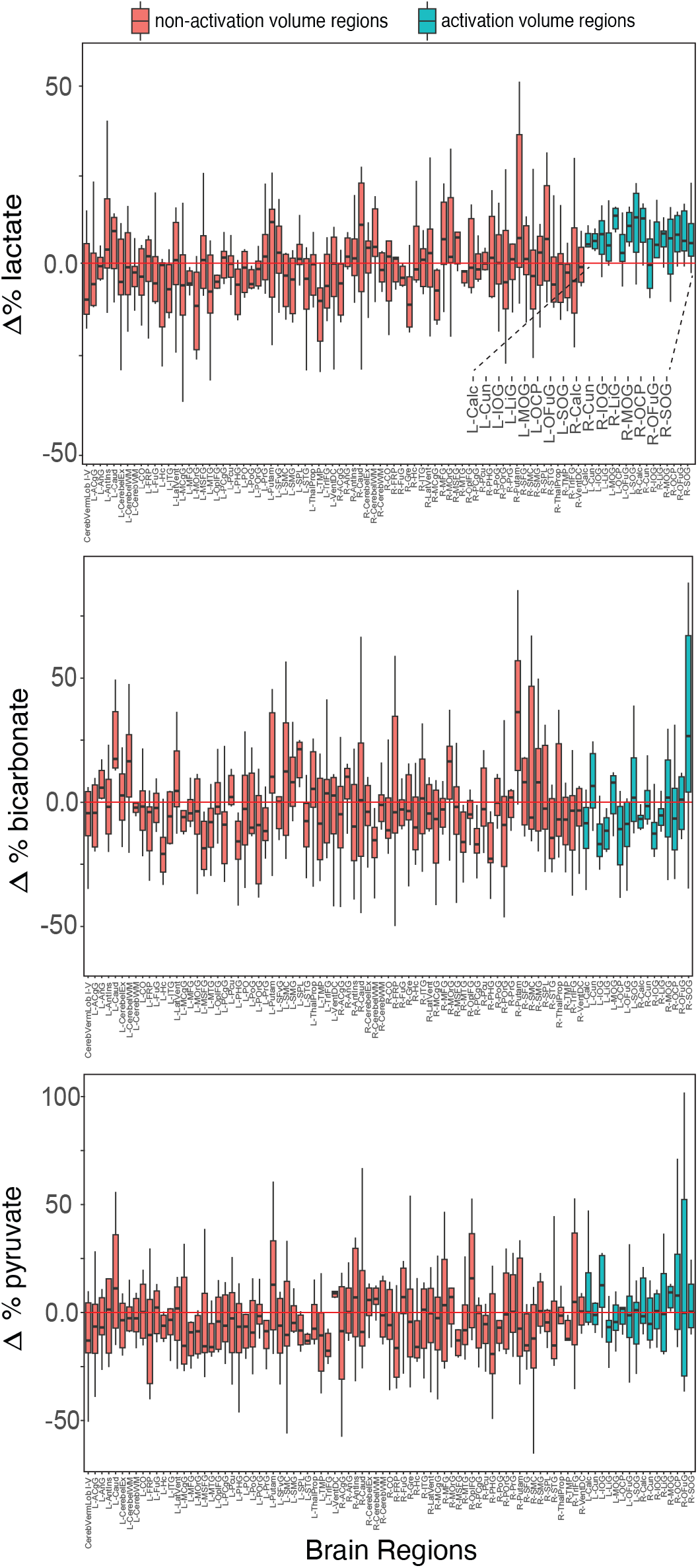
The percentage change between task and control ^13^C-scans for each BrainCOLOR region. The BOLD fMRI t-statistic maps were to designate ‘activation volume regions’ and ‘non-activation volume regions’. A elevated Δ% in ^13^C-lactate is visible in the activation volume regions.

Surprisingly, a task-induced increase in ^13^C-bicarbonate signal was not detected. A prior study using HP-^13^C MR spectroscopy found increased ^13^C-bicarbonate signal in the occipital lobe during functional activation, attributed to increased pyruvate oxidation (31). This discrepancy may be because ^13^C-bicarbonate production is partly derived from oxidative metabolism of ^13^C-lactate, as observed in prior studies (32). Due to the high e effective flip angle used at the ^13^C-lactate frequency, ^13^C-bicarbonate signal resulting from ^13^C-lactate would be suppressed. This is consistent with the results of Zaidi et al., where a reduced flip angle is used for the ^13^C-lactate frequency and an increase in ^13^C-bicarbonate is seen during a similar functional activation stimulus (31).

## Conclusions

HP^13^C MRI was used to assess functional activation in a cohort of healthy volunteers. Increased ^13^C-lactate signal in visual cortex regions, defined by BOLD fMRI activation, was observed following a visual stimulus. Thus HP^13^C MRI may be useful for studying stimulus-induced changes in brain lactate metabolism, a distinct pathway implicated in both neuron and astrocyte bioenergetics. Future work will explore different data acquisition strategies to better understand the underlying physiological origins of the observed ^13^C-lactate signal.

## Abbreviations

HP^13^C: hyperpolarized ^13^C
EPI: echo-planar imaging
MRI: magnetic resonance imaging FOV field of view
PET: positron emission tomography
fMRI: functional magnetic resonance imaging
fMRS: functional magnetic resonance spectroscopy

## Acknowledgments

Funding support from the Canadian Cancer Society grant 707455, Canadian Institutes of Health Research grant PJT-152928 and National Institutes of Health grant R01CA237466. K.R.K. is supported by the National Institutes of Health NIH/NCI Cancer Center Support Grant P30CA008748 and the Center for Molecular Imaging andBioengineering (CMIB) at Memorial Sloan Kettering Cancer Center.

## Author Contributions

H.S. and C.H.C. are co-senior authors; B.U., N.I.C.C., F.T., and C.H.C designed the study protocol; N.M. and W.J.P. prepared pharmacy samples for injection; B.U., N.I.C.C, N.D.B, R.E., and H.S. acquired data; B.U. analyzed data; B.U., A.P., S.J.G., C.H., K.R.K., H.S., and C.H.C. interpreted results; B.U., N.I.C.C., and C.H.C. prepared figures and manuscript; B.U., N.I.C.C., S.J.G, K.R.K., A.P., H.S., and C.H.C. edited and revised manuscript; all co-authors approved final version of manuscript.

## REFERENCES

1. Jan H Ardenkj r-Larsen, Björn Fridlund, Andreas Gram, Georg Hansson, Lennart Hansson, Mathilde H Lerche, Rolf Servin, Mikkel Thaning, and Klaes Golman. Increase in signal-to-noise ratio of> 10,000 times in liquid-state nmr. Proceedings of the National Academy of Sciences, 100(18):10158–10163, 2003.

2. Vesselin Z Miloushev, Kristin L Granlund, Rostislav Boltyanskiy, Serge K Lyashchenko, Lisa M DeAngelis, Ingo K Mellingho, Cameron W Brennan, Vivian Tabar, T Jonathan Yang, Andrei I Holodny, et al. Metabolic imaging of the human brain with hyperpolarized 13c pyruvate demonstrates 13c lactate production in brain tumor patients. Cancer research, 78(14):3755–3760, 2018.

3. James T Grist, Mary A McLean, Frank Riemer, Rolf F Schulte, Surrin S Deen, Fulvio Zaccagna, Ramona Woitek, Charlie J Daniels, Joshua D Kaggie, Tomasz Matyz, et al. Quantifying normal human brain metabolism using hyperpolarized [1 13c] pyruvate and magnetic resonance imaging. NeuroImage, 2019.

4. Jeremy W Gordon, Hsin-Yu Chen, Adam Autry, Ilwoo Park, Mark Van Criekinge, Daniele Mammoli, Eugene Milshteyn, Robert Bok, Duan Xu, Yan Li, et al. Translation of carbon-13 epi for hyperpolarized mr molecular imaging of prostate and brain cancer patients. Magnetic resonance in medicine, 81(4):2702–2709, 2019.

5. Casey Y Lee, Hany Soliman, Benjamin J Geraghty, Albert P Chen, Kim A Connelly, Ruby Endre, William J Perks, Chris Heyn, Sandra E Black, and Charles H Cunningham. Lactate topography of the human brain using hyperpolarized 13c-mri. Neuroimage, 204:116202, 2020.

6. Casey Y Lee, Hany Soliman, Nadia D Bragagnolo, Arjun Sahgal, Benjamin J Geraghty, Albert P Chen, Ruby Endre, William J Perks, Jay S Detsky, Eric Leung, et al. Predicting response to radiotherapy of intracranial metastases with hyperpolarized 13 13 c mri. Journal of Neuro-oncology, 152:551–557, 2021.

7. Biranavan Uthayakumar, Hany Soliman, Nadia D Bragagnolo, Nicole IC Cappelletto, Casey Y Lee, Benjamin Geraghty, Albert P Chen, William J Perks, Nathan Ma, Chris Heyn, et al. Age-associated change in pyruvate metabolism investigated with hyperpolarized 13c-mri of the human brain. Human Brain Mapping, 2023.

8. Fawzi Boumezbeur, Kitt F Petersen, Gary W Cline, Graeme F Mason, Kevin L Behar, Gerald I Shulman, and Douglas L Rothman. The contribution of blood lactate to brain energy metabolism in humans measured by dynamic 13c nuclear magnetic resonance spectroscopy. Journal of Neuroscience, 30(42):13983–13991, 2010.

9. Jens Frahm, Gunnar Krüger, Klaus-Dietmar Merboldt, and Andreas Kleinschmidt. Dynamic uncoupling and recoupling of perfusion and oxidative metabolism during focal brain activation in man. Magnetic resonance in medicine, 35(2):143–148, 1996.

10. Benî t Schaller, Ralf Mekle, Lijing Xin, Nicolas Kunz, and Rolf Gruetter. Net increase of lactate and glutamate concentration in activated human visual cortex detected with magnetic resonance spectroscopy at 7 tesla. Journal of neuroscience research, 91(8):1076–1083, 2013.

11. Silvia Mangia, Ivan Tkáč, Rolf Gruetter, Pierre-Francois Van De Moortele, Federico Giove, Bruno Maraviglia, and K mil Uurbil. Sensitivity of single-voxel 1h-mrs in investigating the metabolism of the activated human visual cortex at 7 t. Magnetic resonance imaging, 24(4):343–348, 2006.

12. Ziad S Nasreddine, Natalie A Phillips, Valérie Bédirian, Simon Charbonneau, Victor Whitehead, Isabelle Collin, Je rey L Cummings, and Howard Chertkow. Montreal cognitive assessment. The American Journal of Geriatric Psychiatry, 2003.

13. Benjamin J Geraghty, Justin YC Lau, Albert P Chen, and Charles H Cunningham. Dual-echo epi sequence for integrated distortion correction in 3d time-resolved hyperpolarized 13c mri. Magnetic Resonance in Medicine, 79(2):643–653, 2018.

14. Mark Jenkinson, Peter Bannister, Michael Brady, and Stephen Smith. Improved optimization for the robust and accurate linear registration and motion correction of brain images. Neuroimage, 17(2):825–841, 2002.

15. Arno Klein and Jason Tourville. 101 labeled brain images and a consistent human cortical labeling protocol. Frontiers in neuroscience, 6:171, 2012.

16. Yuankai Huo, Zhoubing Xu, Yunxi Xiong, Katherine Aboud, Prasanna Parvathaneni, Shunxing Bao, Camilo Bermudez, Susan M Resnick, Laurie E Cutting, and Bennett A Landman. 3d whole brain segmentation using spatially localized atlas network tiles. NeuroImage, 194:105–119, 2019.

17. Bruce Fischl. Freesurfer. Neuroimage, 62(2):774–781, 2012.

18. Somnath Datta and Glen A Satten. A signed-rank test for clustered data. Biometrics, 64(2):501–507, 2008.

19. Robert W Cox. Afni: software for analysis and visualization of functional magnetic resonance neuroimages. Computers and Biomedical research, 29(3):162–173, 1996.

20. Robert W Cox and James S Hyde. Software tools for analysis and visualization of fmri data. NMR in Biomedicine: An International Journal Devoted to the Development and Application of Magnetic Resonance In Vivo, 10(4-5):171–178, 1997.

21. Choong-Wan Woo, Anjali Krishnan, and Tor D Wager. Cluster-extent based thresholding in fmri analyses: pitfalls and recommendations. Neuroimage, 91:412–419, 2014.

22. John C Mazziotta, Arthur W Toga, Alan Evans, Peter Fox, Jack Lancaster, et al. A probabilistic atlas of the human brain: theory and rationale for its development. Neuroimage, 2(2):89–101, 1995.

23. Peter T Fox, Marcus E Raichle, Mark A Mintun, and Carmen Dence. Nonoxidative glucose consumption during focal physiologic neural activity. Science, 241(4864):462–464, 1988.

24. Benoît Schaller, Lijing Xin, Kieran O’Brien, Arthur W Magill, and Rolf Gruetter. Are glutamate and lactate increases ubiquitous to physiological activation? a 1h functional mr spectroscopy study during motor activation in human brain at 7 tesla. Neuroimage, 93:138–145, 2014.

25. Minjie Zhu, Aditya Jhajharia, Sonal Josan, Jae Mo Park, Yi-Fen Yen, Adolf Pfe erbaum, Ralph E Hurd, Daniel M Spielman, and Dirk Mayer. Investigating the origin of the 13c lactate signal in the anesthetized healthy rat brain in vivo after hyperpolarized [1-13c] pyruvate injection. NMR in Biomedicine, page e5073, 2023.

26. Mary C McKenna. The glutamate-glutamine cycle is not stoichiometric: Fates of glutamate in brain. Journal of neuroscience research, 85(15):3347–3358, 2007.

27. Duanghathai Pasanta, Jason L He, Talitha Ford, Georg Oeltzschner, David J Lythgoe, and Nicolaas A Puts. Functional mrs studies of gaba and glutamate/glx a systematic review and meta-analysis. Neuroscience & Biobehavioral Reviews, 144:104940, 2023.

28. Robert G Shulman, Fahmeed Hyder, and Douglas L Rothman. Lactate e ux and the neuroenergetic basis of brain function. NMR in Biomedicine: An International Journal Devoted to the Development and Application of Magnetic Resonance In Vivo, 14(7-8):389–396, 2001.

29. Carlos Manlio Daz-Garca, Rebecca Mongeon, Carolina Lahmann, Dorothy Koveal, Hannah Zucker, and Gary Yellen. Neuronal stimulation triggers neuronal glycolysis and not lactate uptake. Cell metabolism, 26(2):361–374, 2017.

30. Tae Kim, Kristy S Hendrich, Kazuto Masamoto, and Seong-Gi Kim. Arterial versus total blood volume changes during neural activity-induced cerebral blood flow change: implication for bold fmri. Journal of Cerebral Blood Flow & Metabolism, 27(6):1235–1247, 2007.

31. Maheen Zaidi1, Jun Chen, Junjie Ma1, Xiaodong L Wen, Bartnik-Olson Brenda, and Jae M Park. Assessment of human brain pyruvate oxidation using functional hyperpolarized 13c mrs. Proceedings of the Joint Annual ISMRM-ESMRMB 2022, and ISMRT Annual Meeting, London, UK, 2022.

32. Nikolaj B gh, James T Grist, Camilla W Rasmussen, Lotte B Bertelsen, Esben SS Hansen, Jakob U Blicher, Damian J Tyler, and Christo er Laustsen. Lactate saturation limits bicarbonate detection in hyperpolarized 13c-pyruvate mri of the brain. Magnetic Resonance in Medicine, 2022.

